# Brain Modeling ToolKit: an Open Source Software Suite for Multiscale Modeling of Brain Circuits

**DOI:** 10.1101/2020.05.08.084947

**Authors:** Kael Dai, Sergey L. Gratiy, Yazan N. Billeh, Richard Xu, Binghuang Cai, Nicholas Cain, Atle E. Rimehaug, Alexander J. Stasik, Gaute T. Einevoll, Stefan Mihalas, Christof Koch, Anton Arkhipov

## Abstract

Experimental studies in neuroscience are producing data at a rapidly increasing rate, providing exciting opportunities and formidable challenges to existing theoretical and modeling approaches. To turn massive datasets into predictive quantitative frameworks, the field needs software solutions for systematic integration of data into realistic, multiscale models. Here we describe the Brain Modeling ToolKit (BMTK), a software suite for building models and performing simulations at multiple levels of resolution, from biophysically detailed multi-compartmental, to point-neuron, to population-statistical approaches. Leveraging the SONATA file format and existing software such as NEURON, NEST, and others, BMTK offers consistent user experience across multiple levels of resolution. It permits highly sophisticated simulations to be set up with little coding required, thus lowering entry barriers to new users. We illustrate successful applications of BMTK to large-scale simulations of a cortical area. BMTK is an open-source package provided as a resource supporting modeling-based discovery in the community.

## Introduction

Recent emergence of systematic large-scale efforts for comprehensive characterization of brain cell types, their connectivity, and *in vivo* activity (e.g. (Amunts et al., 2016; Bouchard et al., 2016; Hawrylycz et al., 2016; Koch and Jones, 2016; Martin and Chun, 2016; Vogelstein et al., 2016)) is fundamentally reshaping neuroscience research. As the new extremely rich and multimodal data become increasingly available to the community, the need is more urgent than ever to develop sophisticated modeling approaches that could help distill new knowledge from the exuberant complexity of the brain reflected in these datasets (Einevoll et al., 2019). While computational modeling, when combined with theoretical and experimental approaches, clearly has a lot of potential to bridge properties of single cells with brain connectivity, neural activity, and ultimately organism behavior, building such bridges has proven difficult. Some of the greatest barriers are presented by technical challenges of constructing and simulating large and complex biologically-realistic models, integration of different modeling approaches, and systematic sharing of models with the community. New software tools are required to overcome these challenges and enable easy workflows for the new generation of computational models.

One may argue that simulating a huge number of neurons by itself is not a bottleneck any more (Bezaire et al., 2016; Billeh, 2020; Markram et al., 2015), thanks to availability of supercomputers and the very successful software packages that enable complex and highly parallelizable simulations, such as NEURON (Carnevale and Hines, 2006), NEST (Gewaltig and Diesmann, 2007), GENESIS (Bower and Beeman, 1997), MOOSE (Ray and Bhalla, 2008), Brian (Goodman and Brette, 2008), Xolotl (Gorur-Shandilya et al., 2018), and others. However, existing simulation packages traditionally provide a programming environment for users to develop modeling/simulation software code, rather than data-driven interfaces for interactions with model or simulation data. To build sophisticated models, or even to enable efficient simulations, users often need to become experts in the programming environment and languages specific to a simulation package.

Several tools have been recently developed that address some aspects of these challenges, e.g., NeuroConstruct (Gleeson et al., 2007), LFPy (Hagen et al., 2018; Lindén et al., 2014), BioNet (Gratiy et al., 2018), Open Source Brain (Gleeson et al., 2019), HNN (Neymotin et al., 2020), and NetPyNE (Dura-Bernal et al., 2019). These tools do not necessarily provide their own simulation kernel, but instead may rely on an existing simulation engine, such as NEURON, providing a user-friendly interface to this engine. To achieve this, they take advantage of the recent developments of modeling file formats and universal model description languages such as NeuroML (Cannon et al., 2014; Gleeson et al., 2010), PyNN (Davison et al., 2009), NSDF (Ray et al., 2016), and SONATA (Dai et al., 2020). These new developments indicate very welcome signs of progress in necessary software technology, promising improvements to the practice of modeling in neuroscience.

Building upon these trends, we have developed and present here an extensive package for multiscale modeling and simulation, called the Brain Modeling ToolKit (BMTK). While existing tools typically provide an interface to only one simulation engine (for example, NetPyNE (Dura-Bernal et al., 2019) is a powerful interface specifically to the NEURON simulation engine), BMTK has been explicitly developed to furnish interfaces to multiple simulation engines, providing similar user experience in each case. Currently, BMTK supports biophysically detailed, multi-compartmental simulations with NEURON via the BioNet module (Carnevale and Hines, 2006), point-neuron simulations with NEST (Gewaltig and Diesmann, 2007) via the PointNet module, and population-based simulation with diPDE (Cain et al., 2016) via the PopNet module. Through the FilterNet module, BMTK enables filter-based models and simulations, which are often useful, e.g., for providing inputs to simulations of brain networks. Models at all these levels of resolution can be constructed using the BMTK Builder module. With these capabilities, BMTK offers to users a single convenient environment for modeling and simulations across multiple scales and approaches.

From the implementation point of view, BMTK is a Python package that can be installed on a personal computer, a cluster or supercomputer, or in a cloud environment. BMTK provides a Python-based modular environment for model building and simulation, where the model building stage is clearly separated from simulation, as some of the applications leveraging real biological complexity of brain composition and connectivity, like empirically driven placement of synapses, can cause model building to be computationally expensive. It is therefore often useful to build a model once and then load such pre-built models from files for every new simulation. For simulations, BMTK provides a user experience requiring little-to-no programming skills: instead of programming, users simply need to manipulate files as inputs and outputs of simulations. However, advanced users can easily extend BMTK capabilities through their own functions, as BMTK’s open-source Python-based design allows for enhancements in a straightforward manner. In other words, one can use BMTK as a simple interface to harness the power of existing simulation engines without the need for programming, or, alternatively, as a programming environment. The diverse capabilities of BMTK are supported by the modeling file format SONATA (Dai et al., 2020), which is unique in that it provides a complete description of models and simulation inputs/outputs (i.e., various properties of cells, connectivity, and activity), employs highly efficient binary solutions for computationally demanding components of models and simulations, and flexibly supports multiple levels of modeling abstraction. Importantly, SONATA is compatible with the neurophysiology data format NWB (Rubel et al., 2019), which makes it easy for BMTK to interface with experimental data stored as NWB files.

BMTK has been developed with an emphasis on complex and large-scale models and simulations. As such, through its integration with the excellent tools such as NEST and NEURON, it provides a powerful interface permitting very efficient simulations of sophisticated models at multiple scales. This enables easy access to a broad spectrum of computational applications leveraging the new streams of complex information about the brain. However, BMTK also easily supports simpler simulations, including small networks or single-neuron simulations. Overall, the tool is designed for user convenience and flexibility. BMTK is provided freely to the community as an open-source software package (https://alleninstitute.github.io/bmtk/) to facilitate development and simulation of models and support systematic model sharing and reproducibility.

## Results

### BMTK Overview

BMTK is a Python-based software package (originally developed for Python 2.7 and currently supporting Python 3.6+) for creating and simulating neural network models at multiple levels of resolution. It is also an open-source software development kit, allowing users to modify the existing functionality and easily add new extensions or modules. Currently BMTK contains a Builder module for creating models and four simulator modules – BioNet, PointNet, PopNet, and FilterNet – for simulating the models at different levels of granularity (**Fig. 1**).

**Figure 1.**
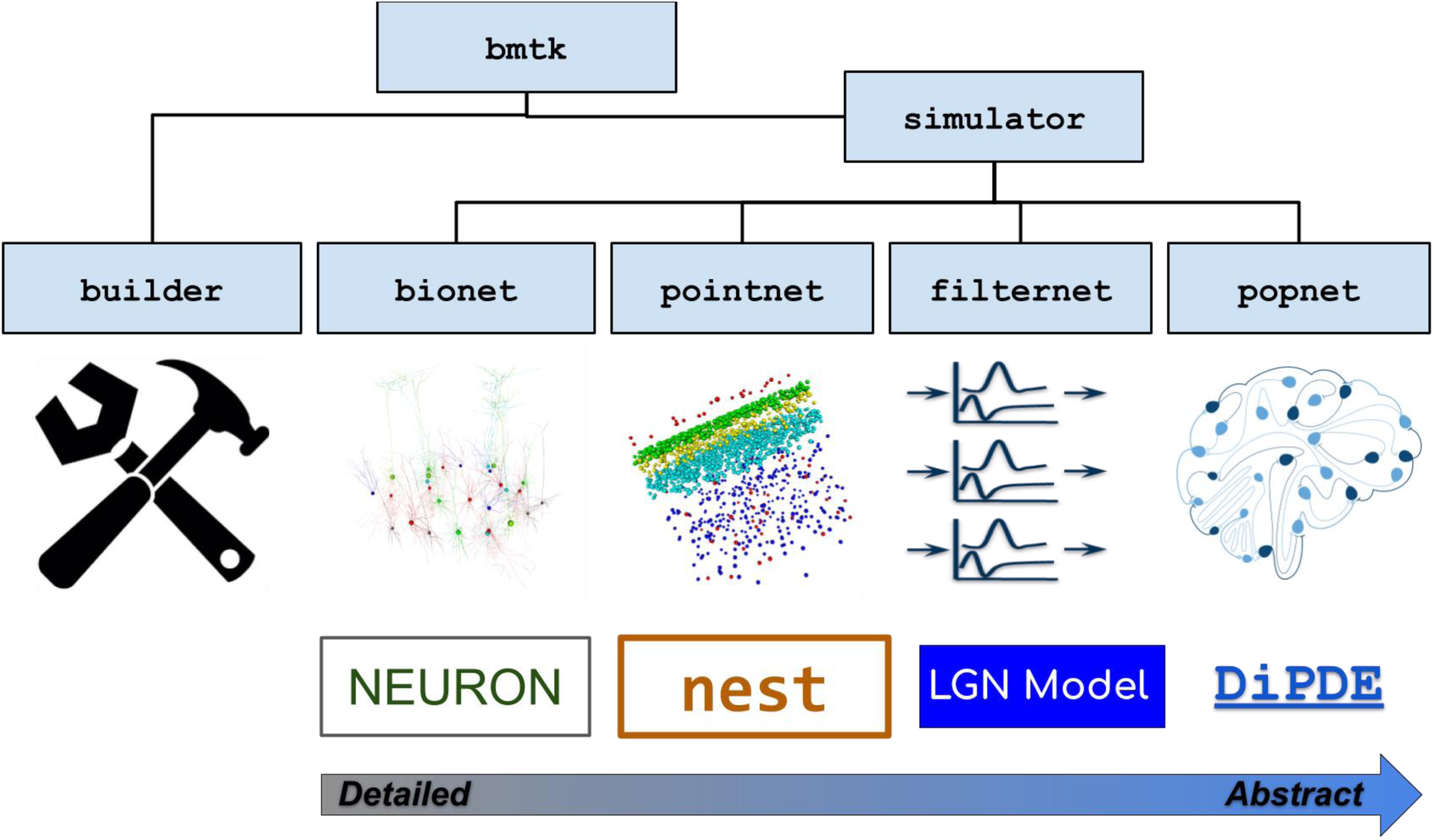
Overview of BMTK. The BMTK software suite consists of several modules. The Builder module contains functions for constructing network models. The simulator modules provide APIs to the simulation engines. BioNet enables simulations of networks consisting of biophysically detailed, multi-compartmental neuron models by interfacing with NEURON. PointNet supports simulations of point-neuron networks via NEST. FilterNet permits simulations of arrays of filters (integrated with the specific case of a model of visual processing by the mouse LGN). PopNet supports simulations with population-statistical models by interfacing with the DiPDE tool. The BMTK modules can subserve multi-stage operations by writing the outputs as files in SONATA format and reading such files as inputs for the next stage of modeling or simulation.

The simulator modules are the application programming interfaces (APIs) to *simulation engines* **(Fig. 1)**, i.e., these modules provide a Python interface to the underlying software packages that execute simulations. The BioNet module provides an interface to NEURON (Carnevale and Hines, 2006) for simulations that involve biophysically detailed, compartmental neuronal models or point-neuron models; PointNet – to NEST (Gewaltig and Diesmann, 2007) for highly efficient point-neuron simulations; PopNet – to the package diPDE (Cain et al., 2016), which implements a population density approach for simulations of coupled networks of neuronal populations; and FilterNet – to BMTK’s built-in solver of filter input-output transformations. The four modules provide a unified user experience for interactions with any of the underlying simulation engines.

Besides the similarity of user experience across modeling levels of resolution, perhaps the main advantage of BMTK to users is that one does not need to become an expert in the programming environments of any of the individual simulation engines, even if one is building and simulating very sophisticated biologically-realistic network models. This is achieved by relying on the standardized data format, SONATA (Dai et al., 2020), for representing model properties and simulation configurations, as well as inputs and outputs. Users only need to provide SONATA files (either by building them using BMTK Builder or by getting files from existing models), and BMTK’s simulator modules will do the rest by translating the SONATA files into model instantiations and simulations by NEURON, NEST, or other engines (**Fig. 2**). Not only does the SONATA format enable this simple workflow under BMTK, it also supports easy model sharing across software packages, as SONATA is implemented in a broad range of modeling tools, such as Blue Brain’s Brion/Brain (https://github.com/BlueBrain/Brion), pyNeuroML (Cannon et al., 2014; Gleeson et al., 2010), pyNN (Davison et al., 2009), and NetPyNE (Dura-Bernal et al., 2019). Moreover, SONATA’s specification for model inputs and output (spikes and time series of membrane voltage, calcium concentration, etc.) is compatible via a converter with the experimental neurophysiology file format NWB (Dai et al., 2020; Rubel et al., 2019).

**Figure 2.**
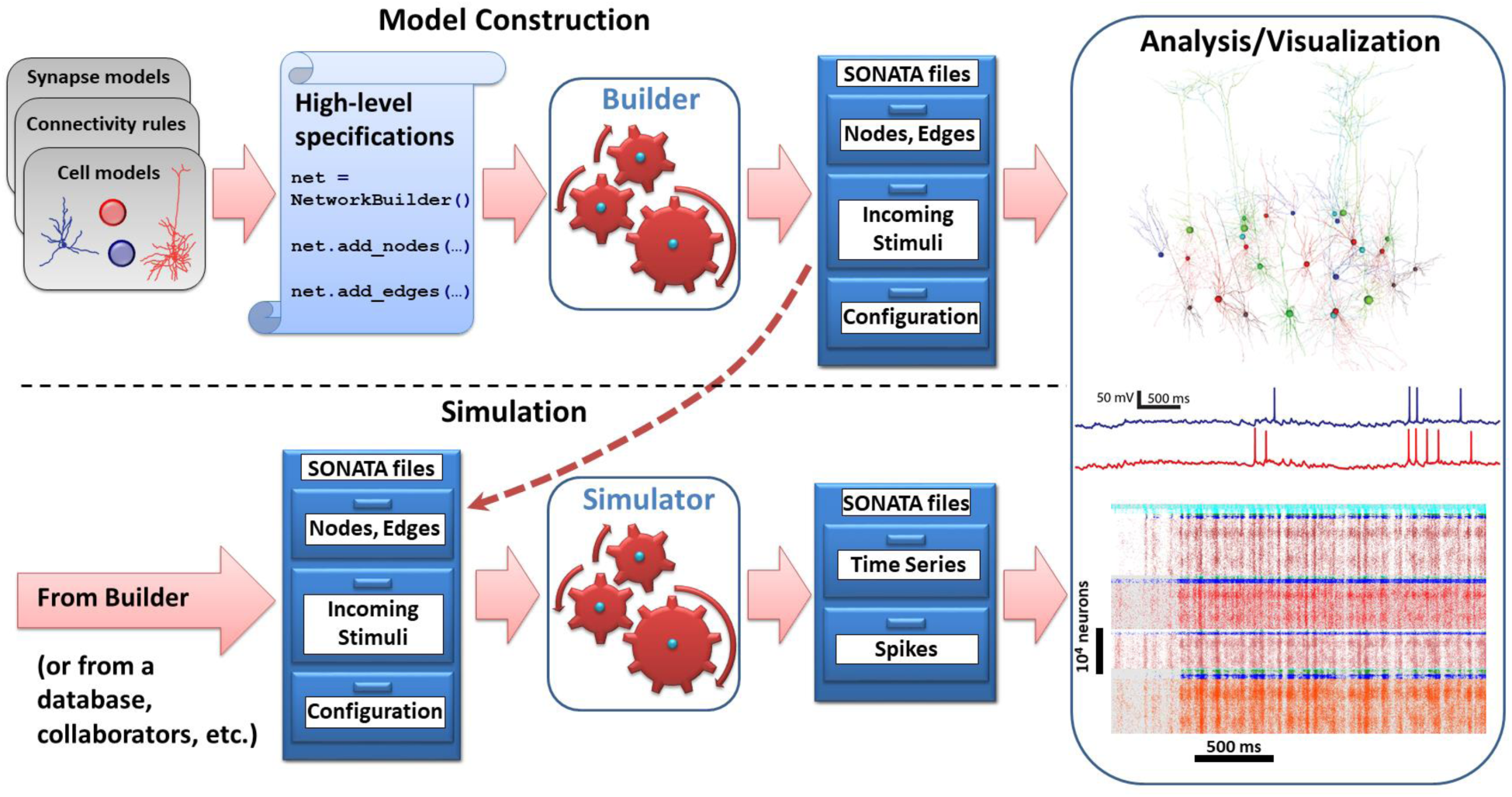
Basic workflow that is conserved across modules of BMTK. Input SONATA files (represented symbolically as chests of drawers) determine the composition and properties of the nodes/network, as well as incoming stimuli (spikes, firing rates, movies) and simulation configuration. Top: the model construction stage. The BMTK Builder combines elements such as cell or synapse models, connectivity rules, and others, via high-level specifications, instantiates the network model, and saves the instantiation as a set of SONATA files. Bottom: simulation stage. The BMTK simulator modules take in the SONATA files as inputs and perform simulations. The input SONATA files may be generated by the BMTK Builder (dashed arrow), any other Builder software supporting SONATA, or from public repositories, collaborators, etc. The BMTK simulator modules produce output, also in SONATA format, typically containing spikes and/or time series (e.g., membrane voltage in selected cells, as a function of time). Right: the SONATA files produced by the BMTK Builder or simulator modules can be analyzed in terms of the model structure or simulated activity (using any analysis software supporting SONATA, or the software that can read HDF5, CSV, and other components of SONATA specification).

As a result, the basic workflow under BMTK is straightforward and consistent across all levels of resolution (**Fig. 2**). Model building is achieved by scripting in Python using the BMTK Builder module, which specify attributes of and relationships between nodes and edges in the constructed network. This step represents the most typical approach currently in use in the modeling field, where descriptive declarations are used to build network instantiations – often constructing very sophisticated networks with only a few lines of code. The output of this module is a set of SONATA files storing model instantiations. The BMTK simulator modules (**Fig. 2**) then run simulations utilizing the SONATA files that describe model composition, inputs (such as incoming spikes), and simulation configuration (duration, etc.). At simulation completion and, if needed, throughout the simulation duration, the simulators write output to disk also in the form of SONATA files.

The BMTK output in SONATA format can be then used for analysis and visualization. Whereas a basic visualization of spiking output or firing rates is provided with BMTK, our design philosophy has been to leave analysis and visualization to other packages. Given that the SONATA format is used for output files and that SONATA can be converted to NWB (Dai et al., 2020; Rubel et al., 2019), analysis of BMTK output is easily achieved with any package that can read SONATA or NWB, or indeed any package that can read the HDF5 format, which underlies both SONATA’s and NWB’s spikes and time series storage. Visualization of the simulated networks can also be achieved with specialized tools as long as they can read SONATA format, which can be easily implemented via the open source pySONATA API (Dai et al., 2020) (https://github.com/AllenInstitute/sonata). One example of such visualization software that reads SONATA is RTNeuron (Hernando et al., 2013), which was used throughout the figures below to visualize examples of BMTK models.

The utility and versatility of BMTK is illustrated below using several examples. First, we describe the BMTK Builder and how it can be used to create simple or very sophisticated network models. Next, we use an example of a simple network consisting of two uniform populations of neurons (excitatory and inhibitory), which we instantiate and simulate using biophysically-detailed compartmental neuronal models in BioNet, point-neuron models in PointNet, and neuronal populations in PopNet. Next, we describe the FilterNet module, which permits one to process stimuli through arrays of filters, currently focusing on converting visual stimuli to spikes that can be used as inputs to simulations of neural networks of vision. Finally, we illustrate the power of BMTK using a variety of real-world applications: simulations of a 230,000-neuron model of mouse V1 implemented at the biophysically detailed and point-neuron levels, computation of the extracellular current source density in simulated cortical tissue, and high-throughput simulations of optogenetic perturbations to diverse cortical cell types.

### Constructing Models with BMTK Builder

The BMTK Builder (**Fig. 3**) is a Python module within the BMTK package. By loading this module, one accesses a variety of functions for building networks and saving results to files in SONATA format. The two major types of tasks performed using the BMTK Builder are instantiating network nodes and instantiating edges.

**Figure 3.**
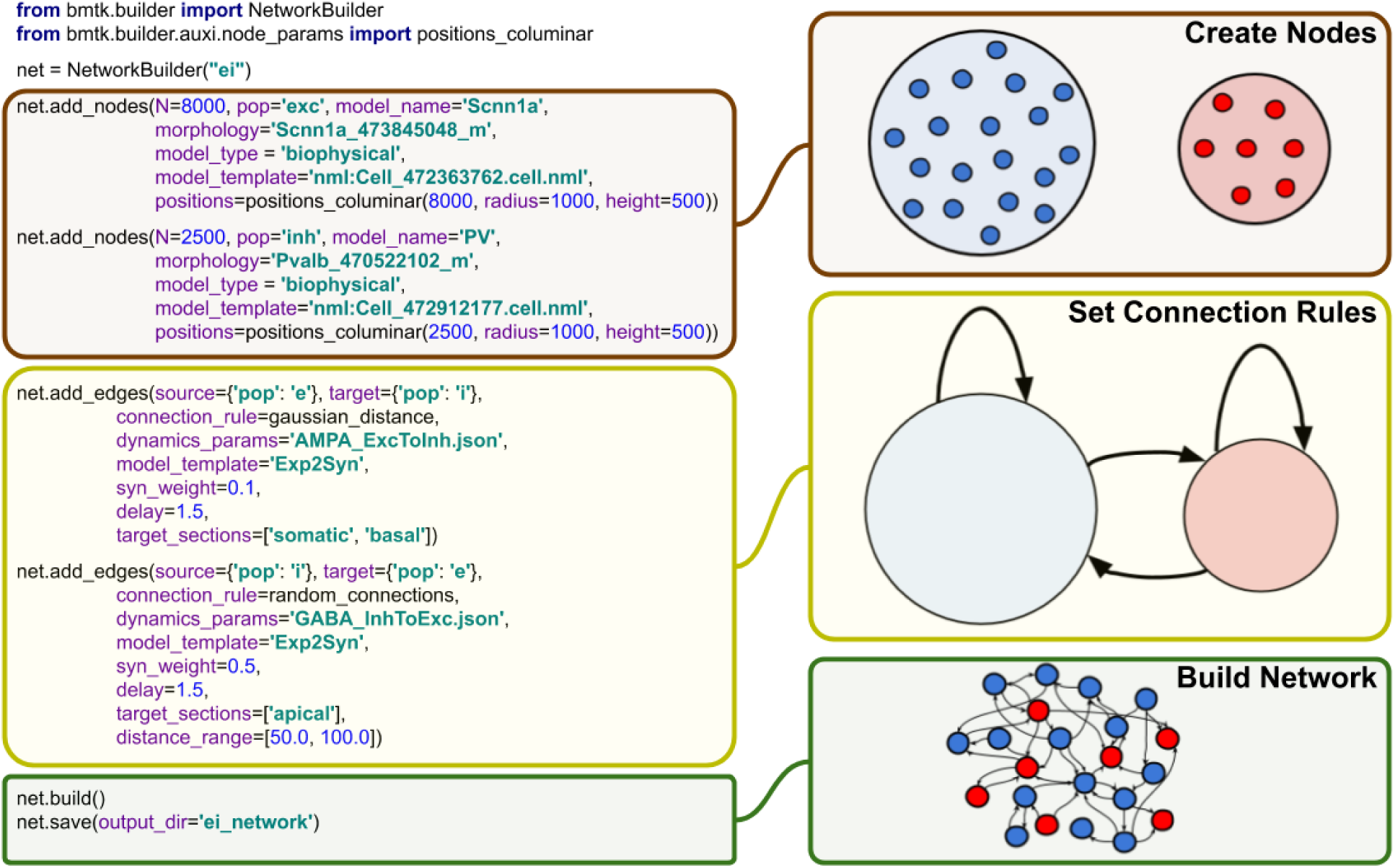
BMTK Builder. The Builder module is used to design and instantiate network models. On the left, examples of the Python commands used in BMTK Builder are presented (simple versions of these commands are shown, for clarity), and on the right purpose of these commands is illustrated schematically on the right. The main stages of model building workflow are defining the nodes and their attributes, defining the connection rules, and then instantiating and saving the network.

When instantiating nodes, one specifies a name for every node type as well as the number of nodes in the type. Furthermore, optional properties of nodes can be specified, such as their positions, types, and other attributes. Some of the attributes are reserved in SONATA format, but otherwise any attributes can be created and assigned as users desire. Functions are provided to distribute values of node properties according to desired distributions (such as distributing cell positions uniformly in a 3D cylindrical volume).

Instantiation of edges follows similar logics. One specifies which populations of nodes should be connected and adds attributes to those connections (edges), some of which are reserved SONATA properties, but otherwise arbitrary attributes can be assigned. BMTK Builder supplies basic functions for establishing probabilistic connectivity between nodes based, for example, on distance between the nodes.

We emphasize that BMTK Builder is designed as a general framework open for extensions. It currently provides functions that, for example, help one to distribute nodes or organize connections according to certain logics, but users are encouraged to utilize their own functions as well. This is easily achieved by the extensible Python interface of the Builder. Additional functions will be added to the core library of the Builder per user feedback.

The BMTK Builder is versatile in that it can create both relatively simple network models or highly complex and biologically realistic network models. Below, we describe simulations of networks illustrating two such cases: a network consisting only of two neuronal populations with random connectivity (Brunel, 2000) and a highly sophisticated network model of mouse V1 consisting of 17 cell classes distributed in space across 6 cortical layers, with multiple connectivity rules that account for cell classes, distances, and tuning of physiological responses (Billeh, 2020). Both networks were prepared using BMTK Builder (for the former model, see examples in https://github.com/AllenInstitute/bmtk, and for the latter, see https://portal.brain-map.org/explore/models/mv1-all-layers). It should be noted that, naturally, complexity of a model, especially of the connectivity rules, strongly influences the computing expense required for model building. For instance, generating the 230,000-neuron V1 model (Billeh, 2020) can take ∼100 CPU-hours or more, depending on the connectivity rules used (note, however, that instantiating a fully actualized model can be parallelized on a cluster). For cases like this, the BMTK’s approach (**Figs. 2, 3**) of building the model and saving it in SONATA files for subsequent simulations, rather than rebuilding the model every time a simulation is run, is clearly beneficial.

A unique feature of BMTK enabled by the SONATA format is that models prepared for one level of resolution can largely be reused for another. For example, a network connectivity created by BMTK Builder for a biophysically detailed simulation contains connections between individual cells as well as descriptions of where synapses should be located on the dendrites of target neurons. This information is stored in SONATA files, which can be used to run a BioNet biophysically detailed simulations. The same files, however, can be used to run a PointNet simulation, which has no representation of dendrites (all neurons are points). In the latter case, only the cell-to-cell connectivity information is used by PointNet, whereas the dendritic locations are ignored. We also note that SONATA files produced by BMTK Builder can be further edited directly, outside of BMTK, since they use well established formats such as HDF5 and CSV (Dai et al., 2020), which can be read and written by many software packages and programming languages.

### Biophysically Detailed, Point-Neuron, and Population Simulations with BioNet, PointNet, and PopNet

For simulating networks of *interacting* nodes, BMTK currently offers support at three levels of resolution: biophysically detailed, compartmental models with BioNet (Gratiy et al., 2018), the interface to NEURON (Carnevale and Hines, 2006); point-neuron models with PointNet, the interface to NEST (Gewaltig and Diesmann, 2007); and population density dynamics models with PopNet, the interface to diPDE (Cain et al., 2016). In all cases, a user provides as an input the SONATA files (Dai et al., 2020) specifying the model (either constructed with BMTK Builder or obtained via other software, such as NetPyNE (Dura-Bernal et al., 2019) or others; **Fig. 2**) and simulation configuration. The latter is supplied in text-based JSON files containing SONATA-compliant specifications of simulation duration, paths to input and output files, etc. (Dai et al., 2020). The BioNet, PointNet, or PopNet will then interpret the files, run the simulation, and provide the output – such as spikes or various time series, e.g., membrane voltage – also in SONATA format. One useful functionality provided by BMTK is writing the output to disk at user-defined intervals during the simulation. In the case of parallelized simulations each CPU core will cache intermediate results produced on the given core, with the final results collated from data across all cores. See Documentation for more details (https://alleninstitute.github.io/bmtk/).

To illustrate applications of BioNet, PointNet, and PopNet, we constructed at each of the three levels of resolution an instance of a simple randomly connected network with 10,000 excitatory neurons and 2,500 inhibitory neurons, receiving excitatory input from 1,000 external neurons (Brunel, 2000) (**Fig. 4**). This network can exhibit a variety of possible dynamical regimes (Brunel, 2000), with different degrees of synchrony and asynchrony between neurons and regularity of spiking of individual neurons. Here we selected one of the possible regimes (the regime with synchronized neuronal populations and regular spiking) for illustration at all three levels of resolution. The implementation of this can be found among the examples at https://github.com/AllenInstitute/bmtk.

**Figure 4.**
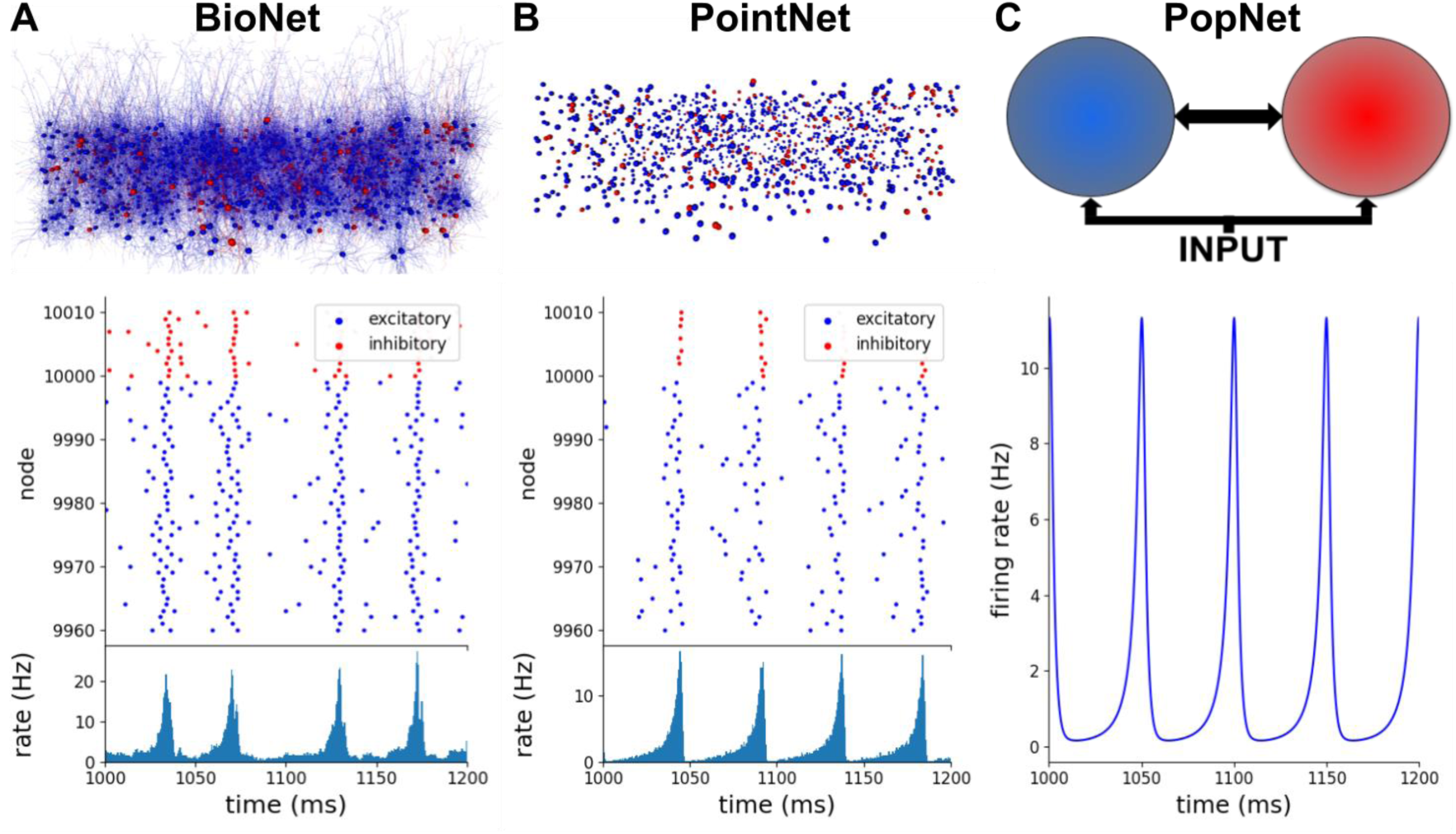
Biophysically detailed, point-neuron, and population simulations with BioNet, PointNet, and PopNet. In all three cases, the interconnected populations of excitatory and inhibitory neurons receive excitatory input from an external population (1,000 Poisson sources firing at the frequency of 150 Hz, replaced by a uniform population in the PopNet case). (A) Biophysically detailed network of randomly connected excitatory and inhibitory neurons, 12,500 total. An RTNeuron visualization of the network is shown alongside its spiking output (spikes from a small portion of the network are shown, for clarity) and the firing rate (for the whole excitatory population) produced by the BMTK’s BioNet module. (B) The same network using the point-neuron approximation. An RTNeuron visualization and simulation output from the BMTK’s PointNet module simulation are shown. (C) Population-based representation of the same network. A schematic of the model and the output of population-density simulation (firing rate for the excitatory population is shown) from BMTK’s PopNet module are illustrated.

We first employed BMTK Builder to construct a 12,500-neuron network model using compartmental neuron representations from the published model of Layer 4 of mouse V1 (Arkhipov et al., 2018), with 264 compartments for each excitatory and 121 compartments for each inhibitory neuron (**Fig. 4A**). The neurons were interconnected with 0.1 probability and received spiking inputs from 1,000 Poisson firing rate sources firing at the frequency of 150 Hz. The model was simulated using BioNet, and we adjusted synaptic parameters to obtain the desired dynamical regime. To compare with the other levels of resolution (below), we plotted the spike rasters and population firing rates, which show that neurons fire in a synchronized and regular fashion (**Fig. 4A**). The population as a whole exhibits the main frequency of ∼20 Hz.

For the PointNet example, we took the model used for the BioNet simulation above and used all of its components applicable to point-neuron simulations – such as the information about which cell connects to which, but not where individual synapses are placed. Naturally, parameters of neurons and of synapses (such as synaptic strengths) needed to be adjusted, as the meaning of many of such parameters are very different between compartmental and point-neuron models. PointNet simulations were carried out, and the synaptic weights were adjusted to obtain the dynamical regime (**Fig. 4B**) similar to that in the BioNet simulation above, with the synchronized neurons emitting bursts of population activity at ∼20 Hz.

Finally, at the PopNet level (**Fig. 4C**), the network was reduced to three nodes – the excitatory, the inhibitory, and the external stimulus populations, with connections between them. After building this very simple network in BMTK Builder, we simulated it with PopNet and adjusted parameters to obtain the desired dynamical regime. Since only the population rate was available here as the output, it was impossible to judge the regularity of firing of individual neurons, but the population activity was clearly similar to the BioNet and PointNet cases. The firing rate exhibited sharp oscillations of population activity at ∼20 Hz, with the activity reaching zero level between each peak, indicating complete silence of all neurons at regular intervals. Note that, like in the BioNet and PointNet cases, the external population here provides a constant level of activity (i.e., individual neurons in the external population fire spikes at irregular intervals according to Poisson statistics, but their collective output at the population level is approximately constant at all times).

### Simulations Using Filter Arrays with FilterNet

Many models of the nervous system utilize filters – mathematical objects that take in multi-dimensional data and return an output, typically by performing a convolution of the input data with certain functions. FilterNet is a module of BMTK that allows users to operate with filters. A typical application may be processing of peripheral sensory input (**Fig. 5**). For example, an array of filters may be used to represent retinal cells, with the input being movies and the output being retinal firing rates or spikes. These output signals in turn can be used as inputs to neurons deeper in the brain explicitly simulated using other modules of BMTK, such as BioNet or PointNet.

**Figure 5.**
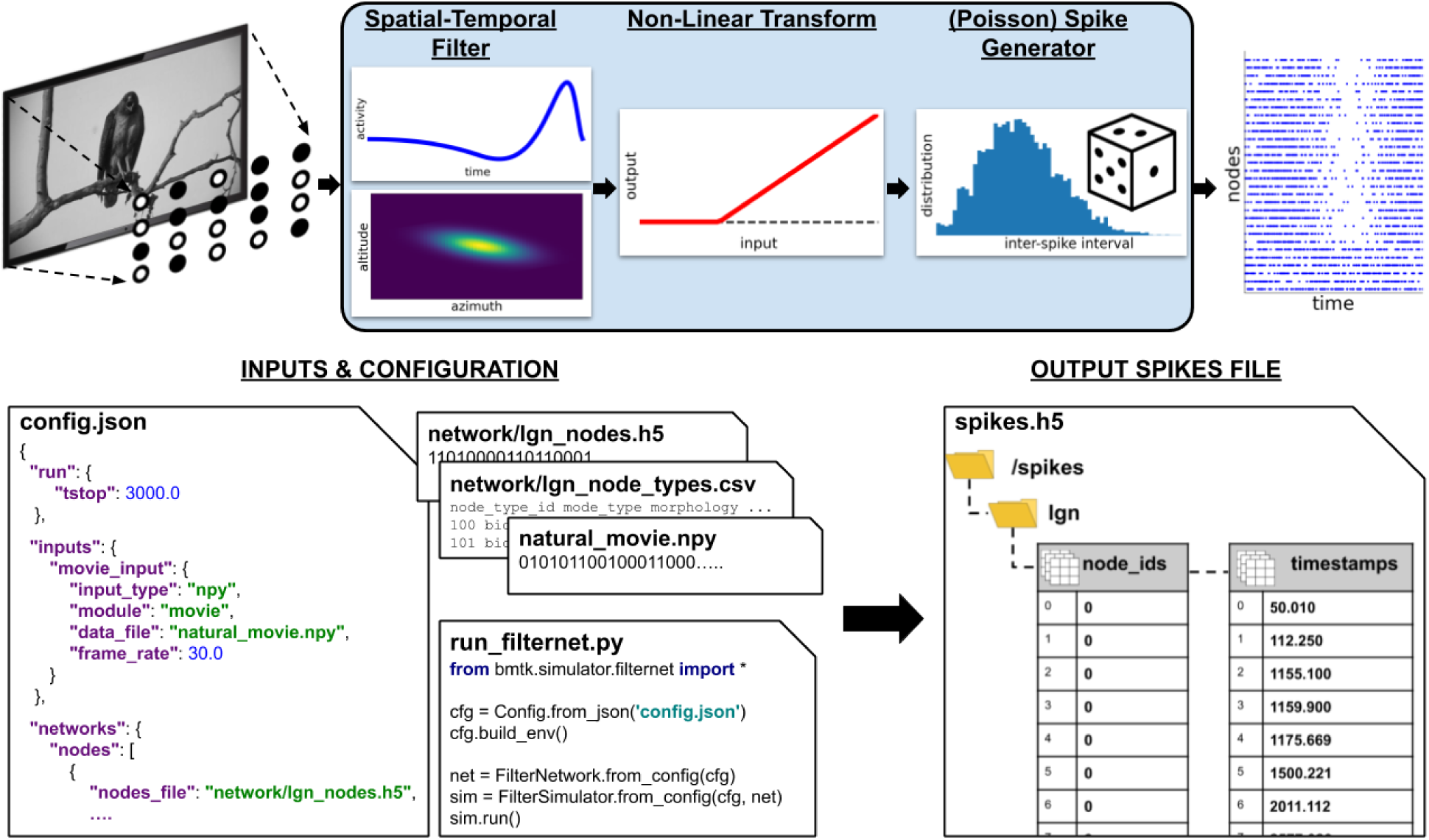
The FilterNet module. Top, general workflow in FilterNet. In case of a visual stimulus, a movie is processed by an array of filters distributed in the visual space. Each filter convolves the frames of the movie with the spatial and temporal kernels, performs rectification, and outputs a time depending firing rate representing the response of the filter to the movie, which can be also converted to instantiations of spike trains. Bottom, illustration of inputs and outputs of FilterNet. Inputs include specifications of parameters such as duration, frame rate, and file locations, as well as contents of the files describing the input patterns and filter properties and distributions. The “run_filternet.py” script is used to carry out the calculations. The output may contain the time series of time-dependent firing rates for each filter and spike trains (illustrated) generated from these time series.

Like the other simulation modules of BMTK, FilterNet is an API that allows users to specify and interact with simulations. FilterNet provides a similar user experience to BioNet, PointNet, and PopNet, in that users work with SONATA-formatted input files that determine functional forms and parameters of the filters, whereas simulation configuration files determine simulation parameters, such as its duration, and location of input and output files.

The current implementation of FilterNet contains the LGNModel simulator, which was created to provide thalamocortical inputs to biologically realistic models of the mouse visual cortex (Arkhipov et al., 2018; Billeh, 2020). This simulator assumes that the input is a movie (a 3D array – two dimensions for space and one for time) and produces the output which is a time-varying firing rate for each filter. A filter here represents an individual cell in the Lateral Geniculate Nucleus (LGN) of mouse thalamus, which projects to the visual cortex. Realistic parameters for such filters, optimized based on the experimental recordings, are available online (http://portal.brain-map.org/explore/models/mv1-all-layers). The FilterNet API can also be easily connected with user-defined functions modeling the input-output filter relationship, which may represent various types of inputs (for example, other sensory stimuli beyond the visual 3D arrays).

An example workflow of FilterNet with LGNModel is illustrated in **Fig. 5**. Here, a movie clip is provided as a 3-dimensional matrix (schematically represented by an image on the top left). A user defines the frame rate, so that the frames can be pinned to the output time axis, and also selects the types of the filters to be used, their numbers, and how they are distributed in the visual space. The types of the filters and their parameters can be taken from our online repository (http://portal.brain-map.org/explore/models/mv1-all-layers) where the filters were optimized to match types of *in vivo* responses of neurons in the mouse LGN (Billeh, 2020; Durand et al., 2016), or one can easily replace these parameters with those of their own choosing. Each filter performs a spatially-temporally separable convolution with the input movie array using two kernels – one operating on the time course of the movie and the other in the visual space (frame pixels). The result of this transformation is rectified. The output of each filter is then a time-varying firing rate, sampled at a frequency defined by the users. FilterNet can also instantiate spike trains from these firing rates using a Poisson process (**Fig. 5**).

In typical applications one runs a simulation where a movie is passed through an array of filters, each filter returning the firing rate and, potentially, a set of instantiated spike trains (each train corresponding to a single trial). These spike trains can be used as inputs to models of neuronal networks (see an example below of a network model of mouse V1 driven by spikes from the LGN, **Fig. 6**). In these applications, the filters become external nodes for other BMTK simulations. Typically, the FilterNet simulations would be done first and their output saved to files, and these outputs would then be reused in subsequent network simulations. The critical intermediate step of determining which filter supplies inputs to which target neuron in the simulated network is accomplished via BMTK Builder, where users can define functions for connecting external nodes to internal ones. The subsequent simulations can be performed with BioNet, PointNet, or PopNet.

**Figure 6.**
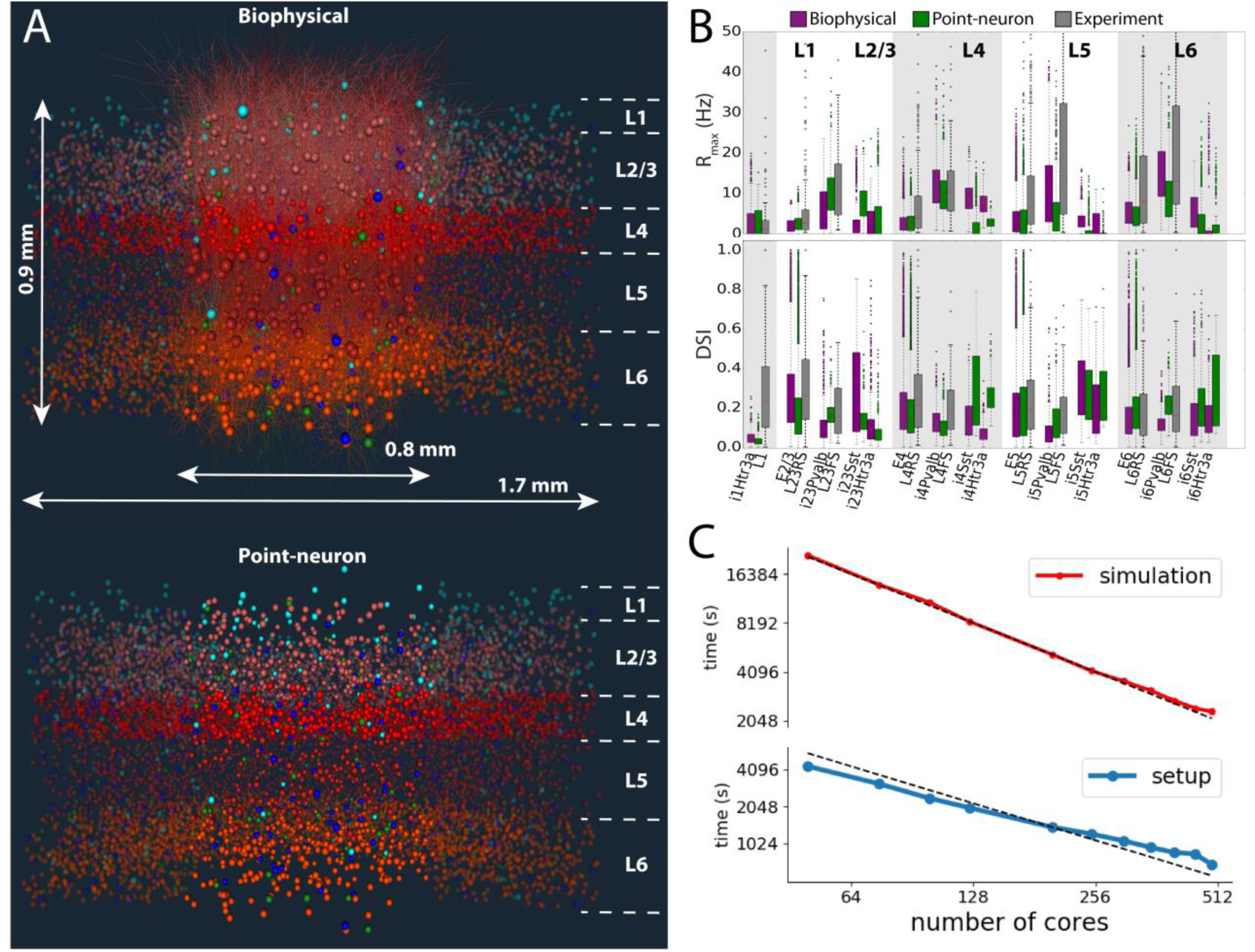
The biophysical and point-neuron V1 models. (A) Visualizations of the biophysical and point-neuron models. The 230,000-neuron models emulate the central portion of the mouse V1, across the full cortical depth, containing layers 1, 2/3, 4, 5, and 6 (layer boundaries are indicated). In the top model, the core portion, ∼50,000 neurons, is simulated using biophysically detailed compartmental neuronal models, and the annulus around the core using leaky integrate-and-fire (LIF) point-neuron models. In the bottom model, both core and the annulus employ the generalized LIF neuronal models. Neurons are colored by cell class: hues of red for excitatory cells in layers 2/3, 4, 5, and 6, and blue, cyan and green for Pvalb, SST, and Htr3a inhibitory class. (B) Summary of firing rates and direction selectivity index (DSI) obtained from the biophysical and point-neuron simulations, vs. experimental extracellular electrophysiology recordings, by cell class. The data were obtained from 2.5-second long presentations of drifting gratings at 8 different directions, 10 trials each. “RS” and “FS” are experimentally determined regular- and fast-spiking cells, roughly corresponding to excitatory and Pvalb inhibitory neurons; the SST and Htr3a neurons could not be identified from experiments. (C) Performance benchmarks and scaling of simulations and setup of the biophysical version of the V1 model using BMTK’s BioNet. The simulation involved 0.5 s presentation of gray screen and 2.5 s of a drifting grating. The time shown is the wallclock time it took to obtain 1 second of simulated time, averaged over 3 s of simulation. The dashed lines indicate ideal scaling (relative to 125 cores, which is a typical choice for simulation of such scale).

### Examples of BMTK Applications to Biological Problems

Finally, we present real-life examples of scientific simulations of brain circuits using BMTK. We illustrate large-scale simulations of highly complex brain networks at different levels of resolution (**Fig. 6**); computation of an extracellular electric potential, which is an observable relating the network activity with measurements of a physical signal (**Fig. 7**); and versatile perturbations of network components to mimic optogenetic experiments (**Fig. 8**).

**Figure 7.**
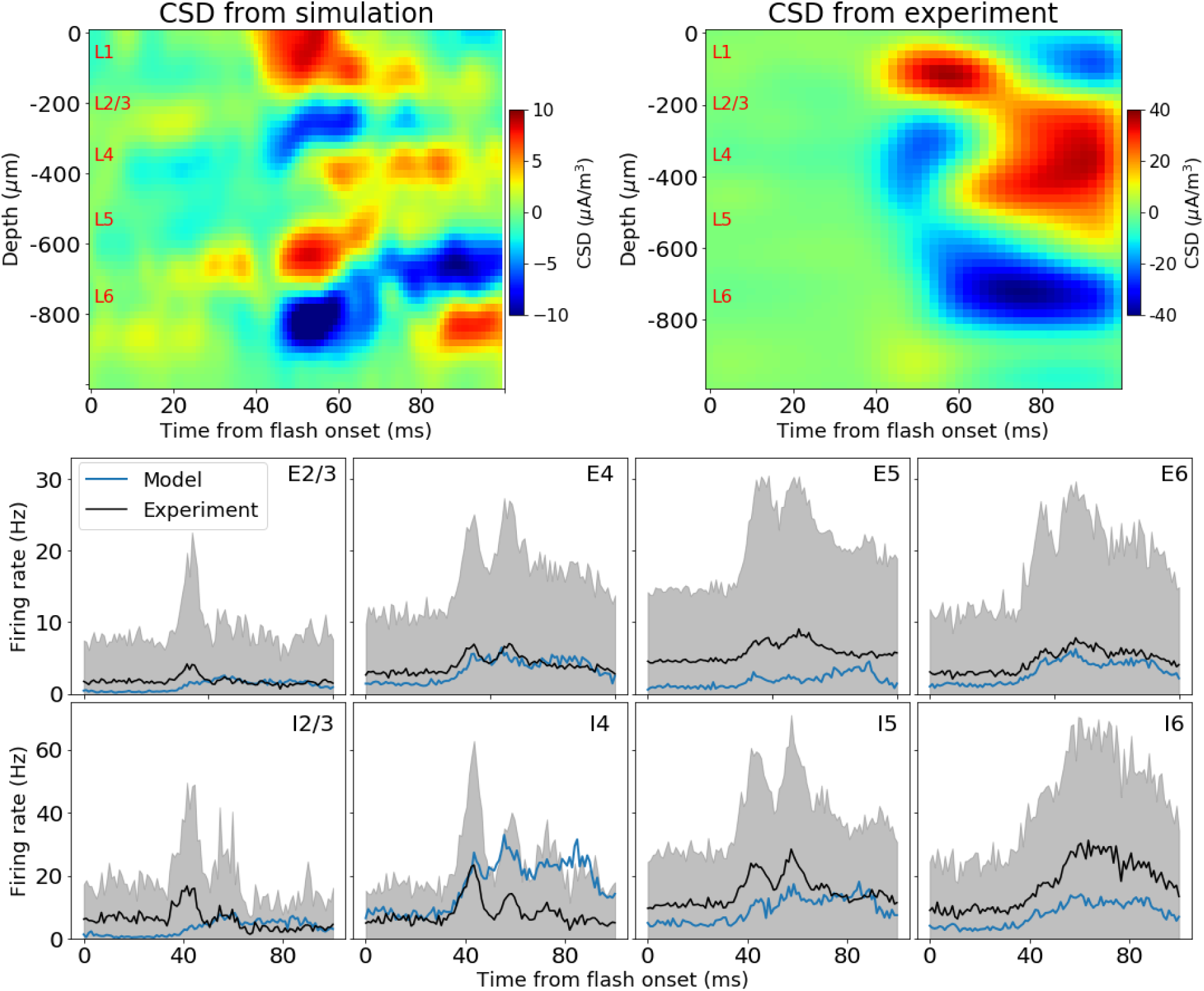
Computing extracellular field potential in BMTK. A simulation using a version of the V1 model (**Fig. 6**) with the full-field flash stimulus is illustrated. The BioNet module of BMTK was used to run the simulation and compute the extracellular potential at multiple virtual electrode locations along the cortical depth; consequently, the potential was used to obtain the Local Field Potential and Current Source Density (CSD). Top: CSD from the simulation and from a single mouse in experiment. Bottom: firing rates for the excitatory (“E”) and inhibitory (“I”) populations in each layer (2/3, 4, 5, and 6). Black: experiment mean. Gray: experiment standard deviation. Blue: simulation mean. Simulation rates are averaged over all neurons in population and 10 trials. Experimental data are averaged over all neurons of the given type recorded from 47 mice, 75 trials each.

**Figure 8.**
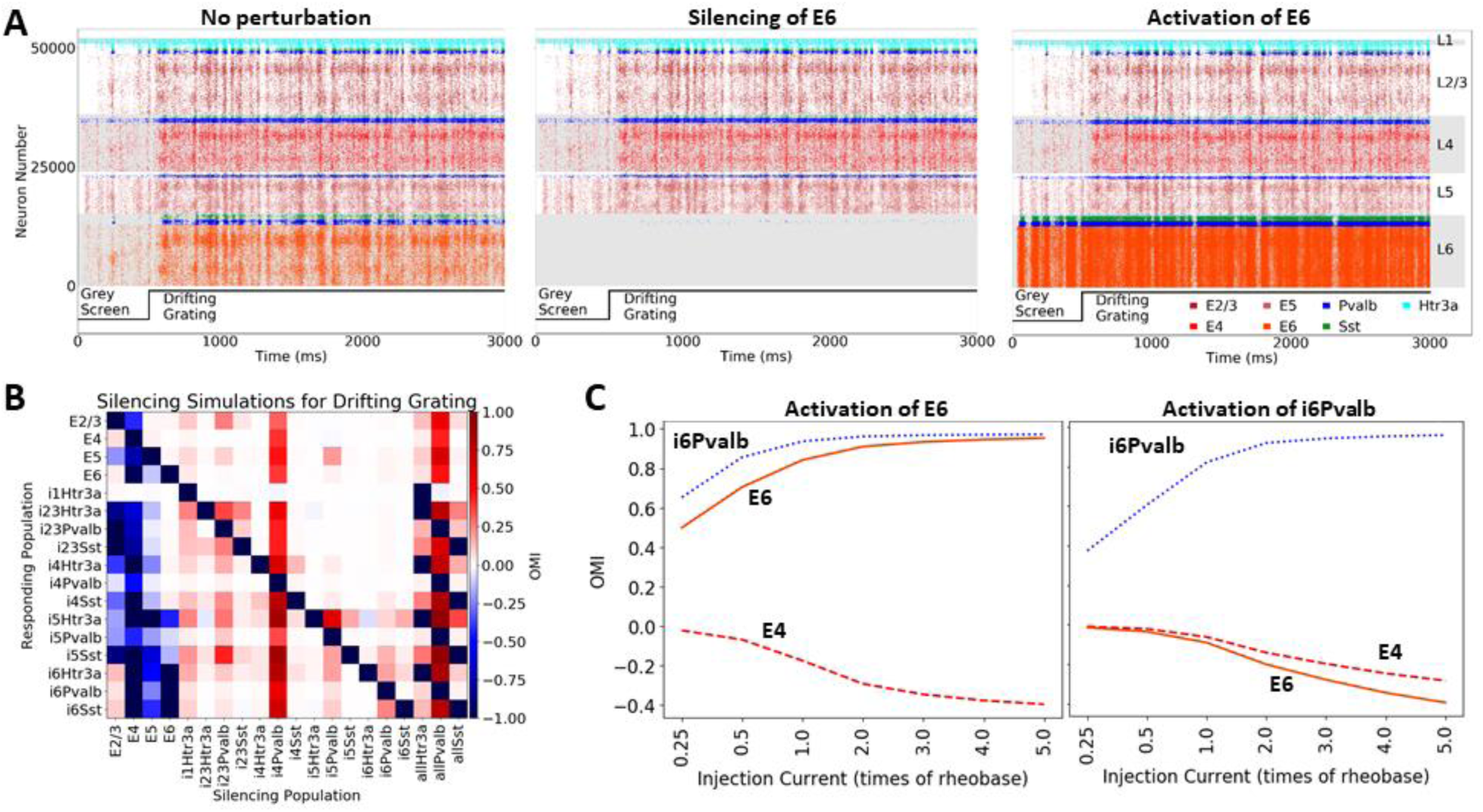
Simulation of optogenetic perturbations using BMTK. The point-neuron version of the V1 model (**Fig. 6**) is used here for illustration. Perturbations are achieved by injecting positive or negative current into cells. (A) Raster plots from 3-second simulations (stimulus: 0.5 s gray followed by 2.5 s of a drifting grating). Simulations without perturbation, with complete silencing of all Layer 6 excitatory cells (E6), and activation of all E6 cells (current equal to 0.5 of the rheobase of each neuron at rest is injected) are illustrated. The perturbation here is applied throughout the course of simulation. (B) Summary of silencing individual cell classes in the V1 model, for the same visual stimulus as in (A). The cell classes listed along the horizontal axis are silenced one by one, and the effect on each cell class (listed along the vertical axis) is characterized using the Optogenetic Modulation Index (OMI; see Main text), averaged over 10 trials and over all cells in the class. The entries “allHtr3a”, “allPvalb”, and “allSst” refer to simulations where, e.g., the Sst class of neurons was silenced in all layers (“allSst”). (C) Activation of Layer 6 excitatory or Pvalb inhibitory neurons, for the same visual stimulus as in (A). Different amplitudes of perturbations are sampled. OMI is computed as in (B), and is shown for 3 select cell classes. Due to inter-laminar projections of Layer 6 Pvalb interneurons to upper layers, activation of either Layer 6 excitatory or Layer 6 inhibitory Pvalb cells leads to the suppression of activity in Layer 4.

### Biophysical and Point-neuron Simulations of the Mouse Cortical Area V1

A recent study (Billeh, 2020) integrated a wide array of experimental information on the composition (cell class, intrinsic properties, and neuron morphologies), connection probabilities and synaptic properties, as well as *in vivo* physiology of neuronal responses in the mouse primary visual cortex (area V1) to construct a comprehensive model of this cortical area (**Fig. 6A**). The model was constructed using the BMTK Builder. It received thalamocortical inputs from the Lateral Geniculate Nucleus (LGN) of the thalamus, which provided the external drive due to visual stimuli (as illustrated in **Fig. 5** for the FIlterNet moduel): 17,400 filters responded to movies (as visual stimuli) and supplied resulting spike trains as inputs to the V1 neurons. These filters represented 14 types of LGN cells, parameterized based on experimental recordings from the LGN (Durand et al., 2016), and were distributed over the whole visual space. The filters were connected to the V1 cells according to experimental data on anatomical and functional properties of the LGN-to-V1 projections (e.g., (Bopp et al., 2017; Ji et al., 2015; Kloc and Maffei, 2014; Lien and Scanziani, 2013, 2018; Morgenstern et al., 2016; Schoonover et al., 2014)). Consequently, arbitrary movies can be used to stimulate the model, enabling direct comparison with experimental trials that used specific movies shown to awake mice while recording extracellular electrophysiology from V1 with the high-throughput Neuropixles probes (Siegle et al., 2019).

The model of V1 was constructed at two levels of resolution: the biophysical level (using compartmental neuron models) and the point-neuron level. The biophysical version was in fact a hybrid model, as the central portion of interest in the model, with ∼50,000 neurons, was represented using compartmental neuron models, whereas the remaining annulus was represented with point-neuron models (**Fig. 6A**). The annulus’s role was primarily to provide a smooth boundary. This hybrid model was simulated with BioNet/NEURON, relying on their ability to handle both compartmental and integrate-and-fire types of models. The fully point-neuron version of the model consisted of Generalized Leaky Integrate-and-Fire (GLIF) neuronal models and was simulated with PointNet/NEST. The neuronal models were sourced from the Allen Cell Types Database (Gouwens et al., 2018, 2019; Teeter et al., 2018).

The two models were each other’s clones, in the sense that they used the same cell positions, individual connections, and all other properties that were applicable to both levels of resolution (as opposed to those applicable to only one level, e.g., dendritic targeting of synapses), the corresponding SONATA files being prepared once in BMTK Builder and then used for both the BioNet and PointNet models. The networks consisted of ∼230,000 neurons, covering all layers of V1 from Layer 1 to Layer 6 and including 17 neuron classes (Billeh, 2020). The models used cell-class-dependent, distance-dependent, and neuron-tuning-dependent connection probability rules and synaptic weight rules. Heavily constrained by experimental data and trained on a small sample of visual stimuli (a single trial of 0.5 s of gray screen and same duration drifting grating), the models generalized well to different stimuli and exhibited many similarities with the experimental recordings. For example, they exhibited firing rates and levels of direction selectivity across cortical layers and cell classes that were similar to experimental ones (**Fig. 6B**). From comparisons of these V1 model simulations to experimental recordings, several predictions were made with regard to the logics of connectivity between cortical cells of different classes, depending on the functional tuning of these cells (Billeh, 2020).

Benchmarks of BioNet simulations of this 230,00-neuron V1 model (**Fig. 6C**) show a close to ideal scaling (i.e., twice faster on twice the number of CPUs) of both the simulation execution time and the model loading time with the number of CPU cores. With the partition of 384 CPU cores, we observe the throughput of approximately 1 second of simulated biological time for slightly over 1 hour of “wall clock” (real) time. These results indicate that extensive simulations for such a large-scale and highly detailed model are possible (Billeh, 2020), although that does require substantial computing resources. On the other hand, we found that the point-neuron version of the V1 model could be simulated efficiently with PointNet on a single CPU core, providing the performance of 1 second of simulated time in approximately 3 minutes of real time. While one gains in speed even further with parallel PointNet simulations of the V1 model, the convenience and speed of the self-contained single-core simulations are such that typically users find them to be the preferred mode for PointNet simulations of such size. Thus, BMTK’s PointNet enables simulations of large-scale models incorporating much biological complexity even with modest computational resources.

It should be noted that the computational performance of BioNet and PointNet relies on the excellent performance and parallelization capabilities of NEURON (Carnevale and Hines, 2006) and NEST (Gewaltig and Diesmann, 2007). What these BMTK modules add is the convenience and interoperability. For example, although NEURON provides powerful parallelization environment, users typically need to write parallel code in that environment to run their simulations. Likewise, constructing sophisticated bio-realistic models in NEURON or NEST requires substantial amount of coding. BMTK streamlines the latter part through the uniform model building operations in BMTK Builder and obviates the former part for the users by dealing with NEURON or NEST parallelization “under the hood”, so that users do not need to write any code at all.

### Computation of the Extracellular Electric Potential

Computing the extracellular field potential in the modeled brain tissue is an important application (Buzsáki et al., 2012; Einevoll et al., 2013, 2019; Gold et al., 2006; Lindén et al., 2011; Mitzdorf, 1987; Senzai et al., 2019) that requires capturing the spatially distributed electric compartments and synapses, as done in biophysically detailed network models. BMTK BioNet’s ability to perform such calculations is illustrated in **Fig. 7**. BioNet allows users to compute the extracellular potential using the line-source approximation (Gratiy et al., 2018; Plonsey, 1974). The potential is then processed to obtain the low-frequency component – the local field potential (LFP), similar to other recently developed tools providing such functionality (e.g., LFPy (Hagen et al., 2018; Lindén et al., 2014), NetPyNE (Dura-Bernal et al., 2019)). BioNet allows users to set up an arbitrary number of recording sites and distribute them in space. One can then use the LFP from multiple electrodes, for example, to compute the current source density (CSD). The resulting LFP and CSD can be directly compared to experimental ones (**Fig. 7**).

The V1 model in **Fig. 6** showed good agreement with experiments for firing rate metrics such as direction selectivity. As a next step, one can use BMTK to investigate the extracellular field dynamics. **Fig. 7** shows one example among a number of model configurations generated (differing, e.g., in the strengths of connections among cell types, the ways how LGN inputs are provided, or distribution of synapses on the neuronal arbors). The CSD and the firing rates across the cortical layers are compared with the experimental data (Siegle et al., 2019). Note that experimental data show substantial variability across mice, and the example from one mouse shown is not representative of all observed CSD patterns. A majority of the 47 mice in this dataset, however, do contain main features seen in **Fig. 7**: an early sink (blue) in Layers 2/3-4 (L2/3-L4), which is then replaced by a source (red), and a delayed but strong sink in L5-L6.

The model captures some of these properties of CSD, though not precisely. The L2/3-4 sink is more sustained than in the experiment, and the later source in these layers is less prominent. The L5-L6 sink starts earlier in the simulation and is narrower along the depth dimension. The overall magnitude of CSD peaks and troughs is also smaller in simulation than in experiment. Nevertheless, it is reassuring that the model captures overall trends in both the dynamics of the firing rates and the major features of CSD (**Fig. 7**). Much further work is necessary to understand how the circuit architecture determines the spiking and LFP/CSD responses. With BMTK and the bio-realistic V1 model (Billeh, 2020), iterations of simulations and adjustments to the model circuit structure will shed light on this question and will lead to improved agreement with experiments.

### Applications to Perturbative Studies of Brain Circuits

BMTK also offers approaches to apply a variety of perturbations and manipulations, which can be specified in the simulation configuration file, e.g., by providing the list of cell IDs to be perturbed and parameterizing the perturbation function. (The scripting interface permits further unlimited possibilities for simulating custom perturbations.) See https://github.com/AllenInstitute/bmtk/blob/develop/docs/tutorial/05_pointnet_modeling.ipynb#5.-Additional-Information

As an example, injection of current directly into neurons is a common technique that can be used effectively to mimic optogenetic perturbations. A follow-up study (Cai et al., 2020) to the V1 model work (Billeh, 2020) used this technique to investigate perturbations of neurons, from single to multiple at a time, selected according to their location, cell class, and functional properties. Many thousands of perturbative simulations were performed using the point-neuron version of the V1 model via the BMTK’s PointNet module. The results agreed with the recent single-neuron optogenetics experiments (Chettih and Harvey, 2019) and suggested coexistence of efficient and robust coding in cortical circuits (Cai et al., 2020). **Fig. 8** shows a complementary set of simulations conducted as part of that project, which consist of silencing or activation of whole cell classes, including titrated perturbations. Currently, BMTK offers an easy way of defining perturbations to either cell populations or a set of individual cells.

**Fig. 8A** shows spiking activity in the core of the V1 model (see **Fig. 6**) in response to visual stimulation with a drifting grating, for a control condition and two types of perturbation to the Layer 6 excitatory cells: complete silencing and modest activation of these neurons. With BMKT, it is easy to sample perturbations to all cell classes in the model and characterize the effect of each on all the other classes. This is illustrated in **Fig. 8B**, which uses the Optogenetic Modulation Index (OMI) to characterize the effect of perturbation. The OMI of a neuron *i* is defined as:

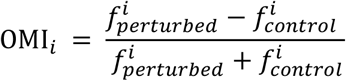

where 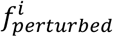 and 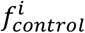 are the firings rate of this neuron during and in the absence of perturbation, respectively. Negative OMI indicates suppression of cell’s firing due to perturbation (OMI = −1 means that the cell is fully suppressed), and positive values indicate elevated firing due to perturbation. Mean OMIs for every cell class in **Fig. 8B** exhibit a rich pattern of various effects depending on the population silenced, including non-intuitive effects of silencing the excitatory populations: e.g., silencing of excitatory populations in Layer 2/3 (E2/3) leads to suppression of E5, but mild activation of E4 and E6.

Furthermore, BMTK permits one to sample the magnitude of perturbation (**Fig. 8C**), which can be done with separate amplitude applied for each cell, e.g., by tying the amount of injected current to the previously measured rheobase of each cell model. **Fig. 8C** shows the effect of such different perturbation magnitudes applied to the excitatory E6 or inhibitory i6Pvalb cell classes. Both perturbations lead to activation of i6Pvalb, but in the first case E6 firing increases, whereas in the second it decreases. Non-intuitively, both perturbations result in suppression of activity in Layer 4. This particular effect of Layer 6 perturbation is due to interlaminar projections from inhibitory Layer 6 Pvalb neurons to upper layers. These results are consistent with the overall inhibitory modulation of superficial layers by Layer 6, demonstrated experimentally (Olsen et al. 2012; Bortone, Olsen, and Scanziani 2014).

Together, these examples demonstrate the capability of BMTK to sample a wide variety of perturbations and therefore enable extensive comparisons with experiments and biologically meaningful predictive studies.

## Discussion

The Brain Modeling ToolKit (BMTK) is a Python package that provides convenient and powerful user interfaces for building and simulating computational models for neuroscience applications. Network models, from very simple to highly complex and biologically realistic, can be constructed using BMTK Builder. BMTK’s FilterNet module provides functionality to process multi-dimensional stimuli via arrays of filters, resulting in time series or spike trains that can be used, e.g., as incoming stimuli for network simulations. The actual network simulations are carried out using BMTK modules BioNet, PointNet, and PopNet, which take advantage of the powerful simulation engines NEURON (Carnevale and Hines, 2006), NEST (Gewaltig and Diesmann, 2007), and diPDE (Cain et al., 2016). Through these modules, BMTK supports simulations at multiple levels of modeling resolution – from filters and population dynamics, to point-neuron and biophysically-detailed compartmental neuronal models.

There are multiple benefits of BMTK for users. The most standard practice in the field is to build relatively simple networks, that can be described by a few lines of code. BMTK is fully compatible with such a practice, as BMTK Builder supports exactly this approach. An additional benefit of modularity is provided by separating the model building and simulating stages, so that it becomes easier to keep track of specific instantiations of models that may be simulated with a variety of different input parameters. On the other hand, a growing area of modeling applications is the development of very sophisticated and biologically realistic models drawing on the extensive experimental datasets, and here BMTK is useful as well. BMTK Builder enables very complex and computationally expensive approaches to constructing network models, as exemplified by the model of mouse V1 described above (Billeh, 2020) (**Fig. 6**). The same example also illustrates how, after constructing a model once, one can reuse many components of the model for simulations at different levels of resolution, such as biophysical with BioNet and point-neuron with PointNet.

Another aspect of benefits to users is the standardization of user experience. The simulation modules of BMTK provide very similar interfaces for interacting with simulations at different levels of resolution, whether with BioNet, PointNet, or PopNet. All steps in the modeling and simulation processes are bound together by employing the SONATA format (Dai et al., 2020) for input and output files. This simplifies and standardizes workflows, and also provides a backbone for sharing models and simulations with the community. Beyond applications in BMTK itself, SONATA ensures a wide spectrum of possibilities for sharing and reusing BMTK models with other tools, and vice versa, since SONATA is supported by or compatible with a growing list of software tools and standards, including NetPyNE, NeuroML, PyNN, RTNeuron, Brion/Brain, and NWB (Cannon et al., 2014; Davison et al., 2009; Dura-Bernal et al., 2019; Gleeson et al., 2010; Hernando et al., 2013; Rubel et al., 2019).

Finally, BMTK enables even non-expert users to perform computationally efficient simulations. The BMTK simulator modules enable simple straightforward simulations, but also harness the excellent capabilities of NEURON (Carnevale and Hines, 2006) and NEST (Gewaltig and Diesmann, 2007) to carry out very large-scale simulations with high computational efficiency, employing parallelization techniques. The latter is an essential requirement for efficient simulations of large and biologically realistic model networks. Previously, in many cases one had to become an expert in parallel programming under the simulator environment and write their own parallel simulation code in that environment. BMTK implements this step for users, so that even users with no experience in programming can perform highly computationally demanding simulations very efficiently. At the same time, due to BMTK’s open-source design as a set of Python modules, those users who are more proficient in software coding can easily implement additional capability of their choice by interfacing their functions with BMTK.

As we showed above, BMTK is a mature tool providing ample opportunities for modeling applications. One can build models, provide realistic inputs, such as visual inputs corresponding to arbitrary movies that might be used in experiments, and perform extensive simulations of brain networks under realistic conditions to obtain a variety of outputs (**Figs. 5, 6**). Current BMTK implementation easily supports output of spikes, membrane voltages, and variables such as calcium concentration. BioNet also permits one to simulate and save the extracellular potential for computing such metrics as LFP and CSD (**Fig. 7**). Importantly, BMTK also permits a variety of perturbations applied to the simulated system, for example in the form of current injections into neurons (**Fig. 8**). One critical application of such capabilities is simulation of optogenetic perturbations of brain circuits, which has become a very powerful tool for interrogating circuit function in experiments (e.g., (Boyden, 2015; Carrillo-Reid et al., 2017; Deisseroth, 2015; Kim et al., 2017; Li et al., 2015, 2019; Madisen et al., 2012)).

BMTK is intended as an open ecosystem that can grow and develop with time. While many useful features are already available based on the initial applications, we intend to add new features, especially driven by user feedback and requests. In addition, BMTK is an open-source project hosted on GitHub (https://alleninstitute.github.io/bmtk/), and users are welcome to submit their own new features and solutions to enhance the tool’s capabilities for everyone’s benefit. We anticipate that BMTK, combined with the SONATA format, can be useful for a broad spectrum of applications on personal computers, supercomputers, and in the cloud environments. Our hope is that BMTK will save effort of many researchers who will be able to focus more on their scientific research and will fuel many discoveries at the interface between modeling, theory, and experimentation.

## Acknowledgments

3-D visualizations were generated using RTNeuron with the support of the Blue Brain Project. We are grateful to Michael Hines for many helpful discussions and suggestions. We wish to thank the Allen Institute founder, Paul G. Allen, for his vision, encouragement, and support.

